# Autoimmunity increases susceptibility to and mortality from sepsis

**DOI:** 10.1101/2021.07.06.451335

**Authors:** Isaac J. Jensen, Samantha N. Jensen, Patrick W. McGonagill, Thomas S. Griffith, Ashutosh K. Mangalam, Vladimir P. Badovinac

## Abstract

Our prior publication detailing how sepsis influences subsequent development of EAE presented a conceptual advance in understanding the post-sepsis chronic immunoparalysis state (Jensen et al., 2020). However, the reverse scenario (autoimmunity prior to sepsis) defines a high-risk patient population whose susceptibility to sepsis remains poorly defined. Herein, we present a retrospective analysis of University of Iowa Hospital and Clinics patients demonstrating increased sepsis incidence among MS, relative to non-MS, patients. To interrogate how autoimmune disease influences host susceptibility to sepsis well-established murine models of MS and sepsis, EAE and CLP, respectively, were utilized. EAE, relative to non-EAE, mice were highly susceptible to sepsis-induced mortality with elevated cytokine storms. These results were further recapitulated in LPS and *S. pneumoniae* sepsis models. This work highlights both the relevance of identifying highly susceptible patient populations and expands the growing body of literature that host immune status at the time of septic insult is a potent mortality determinant.

## Introduction

Multiple sclerosis (MS) is an autoimmune demyelinating disease of the central nervous system (CNS) that affects ~2.8 million individuals worldwide, and cases are rising (Fox, 2004; Walton et al., 2020). The symptomology of MS includes (but is not limited to) pain, motor dysfunction, and cognitive dysfunction. The etiology of MS is not well understood, but is thought to stem from a complex interaction of genetic and environmental factors (Dendrou et al., 2015; Freedman et al., 2018). MS is commonly diagnosed between the ages of 20-40, although underlying subclinical pathogenesis may be present long before diagnosis. MS pathogenesis is mediated by proinflammatory auto-reactive T cells and other immune cells activated prior to migration into the CNS to promote axonal damage (Fox, 2004). In an attempt to subvert the aberrant immune response to the CNS, immunomodulatory/immunosuppressive drugs are often prescribed to patients with MS with varying degrees of success (Tintore et al., 2019). Unfortunately, the use of disease-modifying drugs in patients with MS often comes with increased risk of opportunistic infection (Yong and Kim, 2020). The increased propensity to infection may leave MS patients at an increased risk of sepsis.

Sepsis, a dysregulated host response to infection, impacts 9 people every 6 seconds of which 2 will succumb to the associated cytokine storm (Rudd et al., 2020). Additionally, those who survive demonstrate increased susceptibility to subsequent infection or cancer development (Danahy et al., 2019; Hotchkiss et al., 2016; Jensen et al., 2018a; Jensen et al., 2018b). This increased risk for secondary complication leads to a substantial economic burden costing over $20 billion annually in the United States alone (CDC, 2020). While mortality due to the cytokine storm has diminished over time due to early intervention, the sepsis mortality rate of ~20% is still excessive (Dombrovskiy et al., 2007; Gaieski et al., 2013). Mortality from sepsis is in part due the complexity and interconnectedness of the cytokine storm that is composed of both pro- and anti-inflammatory cytokines (Danahy et al., 2016; Delano and Ward, 2016; Jensen et al., 2021), and is further complicated by individual comorbidities (Rhee et al., 2017; Rhee et al., 2019). The underlying link between MS and subsequent sepsis is not clear. MS patients are often prescribed one of several immunosuppressant drugs, putting them at greater risk of infection. Indeed, certain disease-modifying therapies for MS pose a greater risk for infection, such as rituximab, compared to others (Luna et al., 2020).

Patients with autoimmune diseases, such as MS, are often treated with immunomodulatory drugs that may increase their susceptibility to infection and sepsis. For example, urinary tract infection (UTI) and respiratory infection, are both a common causes of sepsis (Jeganathan et al., 2017) and complications for MS patients, relative to the general population (Harding et al., 2020; Medeiros Junior et al., 2020). In fact, compared to the general healthy population, individuals with MS are at greater risk of sepsis, sepsis-induced complications, and death due to infection (Capkun et al., 2015). MS patients are also more likely to have a principal diagnosis of infection at their final hospital stay prior to death compared to the general healthy population and individuals with diabetes mellitus (Ernst et al., 2016). Moreover, sepsis was a secondary diagnosis for 51% of MS patients compared to 36% and 31% of diabetes mellitus and general healthy individuals, respectively, during a hospital stay (Ernst et al., 2016) demonstrating that even among autoimmune disease MS patients are at increased risk of developing sepsis. The increased propensity to become septic also extends to military veterans, a population that is skewed toward individuals >50 years of age and male (Livingston, 2016), both of which are associated with an increased incidence of sepsis. Lastly, veterans with MS are more likely to be hospitalized and die from infection compared with veterans without MS (Nelson et al., 2015).

We previously studied the impact of sepsis on subsequent MS-like disease using the experimental autoimmune encephalomyelitis (EAE) animal model as a means of conceptually interrogating the immunoparalysis state that occurs after sepsis (Jensen et al., 2020). However, there is a strong need to understand how underlying autoimmune conditions, such as MS, influence susceptibility to sepsis-induced mortality given the increased incidence in this potentially vulnerable population. Thus, with this *Research Advance* we affirm the increased incidence of sepsis in MS patient cohorts relative to non-MS patient cohorts and interrogate how autoimmunity as a comorbidity in septic populations influences susceptibility to sepsis-induced mortality utilizing murine models of MS (EAE) sepsis (cecal ligation and puncture [CLP], LPS, and *S. pneumoniae*).

## Results and Discussion

### MS patients are more prone to sepsis than the general population

Prior literature suggests an increased susceptibility of MS patients to develop sepsis relative to non-MS patient cohorts (Capkun et al., 2015). Therefore, to begin interrogating this potentially interesting interplay, we performed a retrospective analysis of ICU admissions at the University of Iowa Hospital and Clinics. This analysis included 211,470 patients admitted between 2008 and 2020, of which there were 22,930 that were septic and 1,180 that had MS (**Table 1**). Notable features of these patient cohorts included: septic patients tended to be older and male – known risk factors associated with developing sepsis (Rhee et al., 2017; Rhee et al., 2019), while MS patients tended to be female – MS is a known female biased disease (Fox, 2004). There was also a slight increase in the proportion of Caucasian patients among the septic patients. Importantly, MS patients exhibited a significant increase in sepsis incidence (14.4%) relative to non-MS patients (10.8%; Odds ratio: 1.387, p=0.0001). Additionally, while MS patients tended to be female, there was a higher proportion of males among the septic MS patients (35%) relative to the non-septic MS patients (26%). Further, septic MS patients also tended to be older (64+/−14 years) than their non-septic MS patient counterparts (56+/−16 years). These data reaffirm both the higher incidence of sepsis in males and with age even within the MS patient cohort. Overall, these data affirm that MS patients have an increased incidence of sepsis relative to non-MS patient cohorts.

**Table 1.**

| Patient Cohort | All Inpatient | Non-septic | Septic | % Septic | Non-Septic vs Septic |
| --- | --- | --- | --- | --- | --- |
| Total | 211470 | 188540 | 22930 | 10.8% |  |
| Age (+/- SD) | 58 (+/-20) | 57 (+/-21) | 64 (+/-17) |  | <0.0001 |
| % Male | 47% | 46% | 53% |  | <0.0001 |
| % Caucasian | 87% | 86% | 88% |  | <0.0001 |
| Non-MS | 210290 | 187530 | 22760 | 10.8% |  |
| Age (+/- SD) |  | 57 (+/-21) | 64 (+/-17) |  | <0.0001 |
| % Male |  | 46% | 53% |  | <0.0001 |
| % Caucasian |  | 86% | 89% |  | <0.0001 |
| MS | 1180 | 1010 | 170 | 14.4% |  |
| Age (+/- SD) |  | 56 (+/-16) | 64 (+/-14) |  | <0.0001 |
| % Male |  | 26% | 35% |  | 0.0159 |
| % Caucasian |  | 89% | 88% |  | ns |

|  |  | Age (+/-SD) |  | % Male |  | % Caucasian |  |
| --- | --- | --- | --- | --- | --- | --- | --- |
| Non-MS vs MS | Sepsis odds ratio | Non-MS | MS | Non-MS | MS | Non-MS | MS |
|  | 1.387 | 60.5 (+/-5) | 60 (+/-6) | 47% | 27% | 86% | 89% |
| p-value | 0.0001 | ns |  | <0.0001 |  | 0.0121 |  |

### EAE increases host susceptibility to sepsis induce mortality

Given that MS patients have a higher incidence of sepsis, we sought to understand how having an ongoing autoimmune disease would influence host susceptibility to sepsis. To address this relationship, well-established models of inducible MS-like disease and polymicrobial sepsis, EAE and CLP, respectively, were used. C57BL/6 mice were immunized with MOG_35-55_ to induce EAE or left unimmunized (non-EAE). CLP or sham surgery was performed >35 days post-immunization and mortality was assessed (**Figure 1a**). To ensure that mortality was not simply due to ongoing EAE disease, EAE mice were segregated into sham and CLP groups to establish a similar distribution of EAE clinical scores prior to surgery (**Figure 1b**). Non-EAE mice exhibited some mortality, however, EAE mice had diminished survival relative to non-EAE mice (**Figure 1c**). Importantly, EAE mice that underwent sham surgery did not have any mortality, consistent with the model system and demonstrating that mortality in EAE with CLP was not due to EAE disease. These data also suggest the presence of CNS autoimmunity increases the host susceptibility to a fatal septic event. Interestingly, there was an observed relationship between the EAE disease score prior to sepsis induction and the likelihood of mortality (**Figure 1d,e**). Mice with a score of ≤2 had a similar survival rate to naïve CLP mice, whereas all mice with an EAE score >2 succumbed to disease (**Figure 1e**).

**Figure 1:**
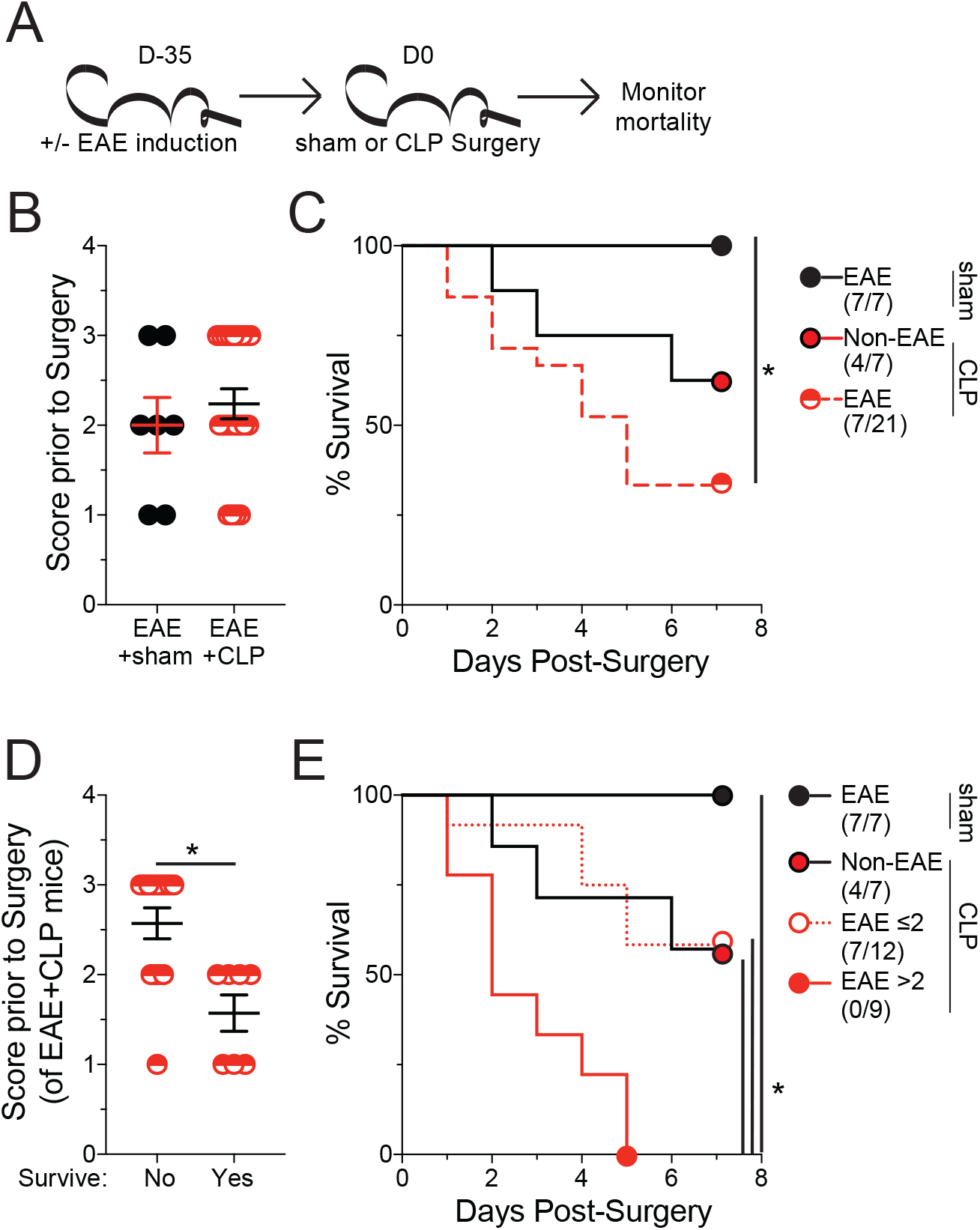
EAE mice have increased susceptibility to sepsis-induced mortality. **A)** Experimental Design: C57BL/6 mice were immunized with MOG_35-55_ to induce EAE. EAE mice underwent either sham or CLP 35 days after EAE induction followed by assessment of mortality, age matched non-immunized (non-EAE) underwent CLP surgery at the same time. **(B)** EAE clinical scores of mice prior to either sham or CLP surgery. **(C)** Kaplan-Meier survival curves of EAE mice that underwent sham (black closed circle) or CLP (red semi-circle) surgery and non-EAE mice that underwent CLP surgery (red closed circle with black outline). **(D)** EAE clinical scores prior to surgery of EAE mice that either succumbed to or survived the septic insult. **(E)** Kaplan-Meier survival curves of EAE mice that underwent sham (black circle), had an EAE score ≤2 prior to CLP (white circle with red outline), or had an EAE score >2 prior to CLP (red closed circle with red outline) surgery and non-EAE mice that underwent CLP surgery (red closed circle with black outline). Data are cumulative of 2 independent experiments with 7-21 mice per group. Error bars represent standard error of the mean. *=p-value<0.05.

### Auto-immune inflammation, not clinical disease, dictates susceptibility to sepsis

The relationship between disease severity and mortality suggests that either the paralysis and associated neurologic damage during EAE is promoting sepsis-induced mortality or differences in the inflammatory response may increase the likelihood of mortality. Indeed, we previously reported that microbially-experienced ‘dirty’ mice with a high degree of immunologic experience are highly susceptible to sepsis-induced mortality due (in part) to elevations in plasma cytokine concentrations both at a baseline and during the peak (~12hrs post-induction) of the cytokine storm (Huggins et al., 2019). Similarly, we have also described a relationship between tumor size at the time of sepsis induction and host mortality (Danahy et al., 2019). Thus, to begin teasing apart the roles of the interconnected phenomena of inflammation and paralysis, mice were immunized at varying times leading up to sepsis induction. This approach establishes a scenario in which disease is subclinical (D5), being established (D15), or fulminant (D25) with ongoing inflammation anticipated in all cohorts (**Figure 2a**). Clinical disease progression occurred in agreement with these expectations (**Figure 2b**). All EAE cohorts, however, exhibited profound susceptibility to sepsis-induced mortality, demonstrating that clinical disease and paralysis were not required for sepsis-induced mortality (**Figure 2c**).

**Figure 2:**
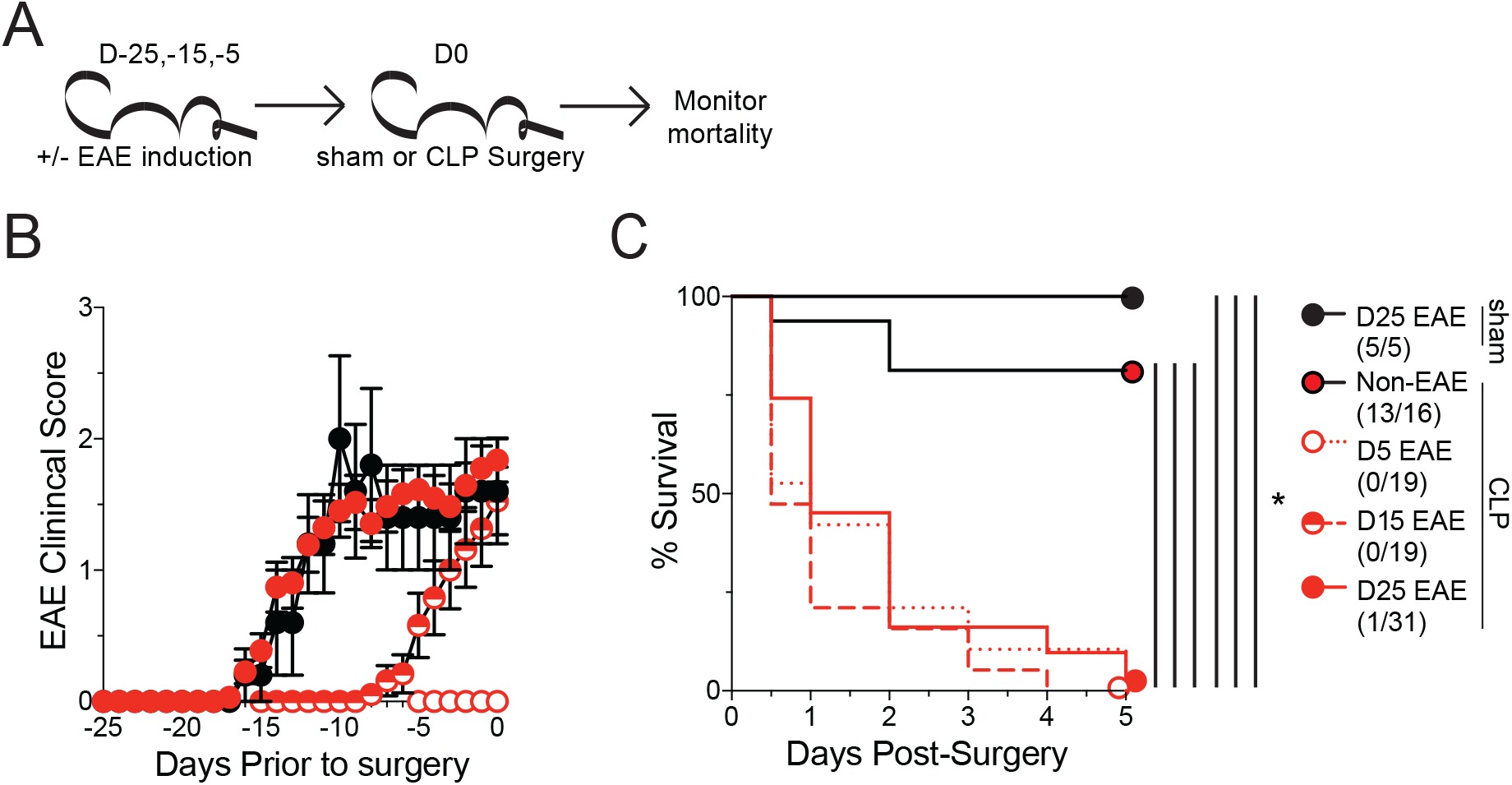
Increased susceptibility of EAE mice to sepsis is independent of disease onset. **A)** Experimental Design: C57BL/6 mice were immunized with MOG_35-55_ to induce EAE at day −25, −15, or −5 prior to either sham or CLP surgery, age matched non-immunized (non-EAE) underwent CLP surgery at the same time. Mortality was monitored in all cohorts. **(B)** EAE clinical scores of mice that were induced for EAE at −25, −15, −5 prior to either sham or sepsis surgery. **(C)** Kaplan-Meier survival curves of day - 25 EAE mice that underwent sham surgery (black circle), non-EAE mice that underwent sepsis surgery (red circle with black outline), and day −25 (red circle with red outline), day −15 (red semi-circle), and day-5 (white circle with red outline) EAE mice that underwent CLP. Data are cumulative of 2 independent experiments with 5-31 mice per group. Error bars represent standard error of the mean. *=p-value<0.05.

To then address the extent to which EAE, similar to infection and cancer, was altering the severity of the sepsis-induced cytokine storm, plasma was collected prior to and 12hrs post-CLP surgery in D5, D15, and D25 (as well as non-EAE) mice and assessed for IL-6, TNF, IL-1β, IFNγ, IL-10, IL2, and IL-12p70 (**Figure 3a**). Importantly, while there was a cytokine storm in all CLP cohorts, the magnitude of the cytokine storm was substantially higher in EAE mice relative to the non-EAE mice (**Figure 3b-d**). Further, EAE mice had a higher baseline expression of many cytokines (**Figure 3c,d**) recapitulating observations in ‘dirty’ mice (Huggins et al., 2019). Of particular note was IL-6 which has previously been described as a strong indicator of the severity of the cytokine storm (Ma et al., 2016; Qiao et al., 2018; Qiu et al., 2018) and was strongly increased in all EAE groups both prior to and after CLP (**Figure 3d**).

**Figure 3:**
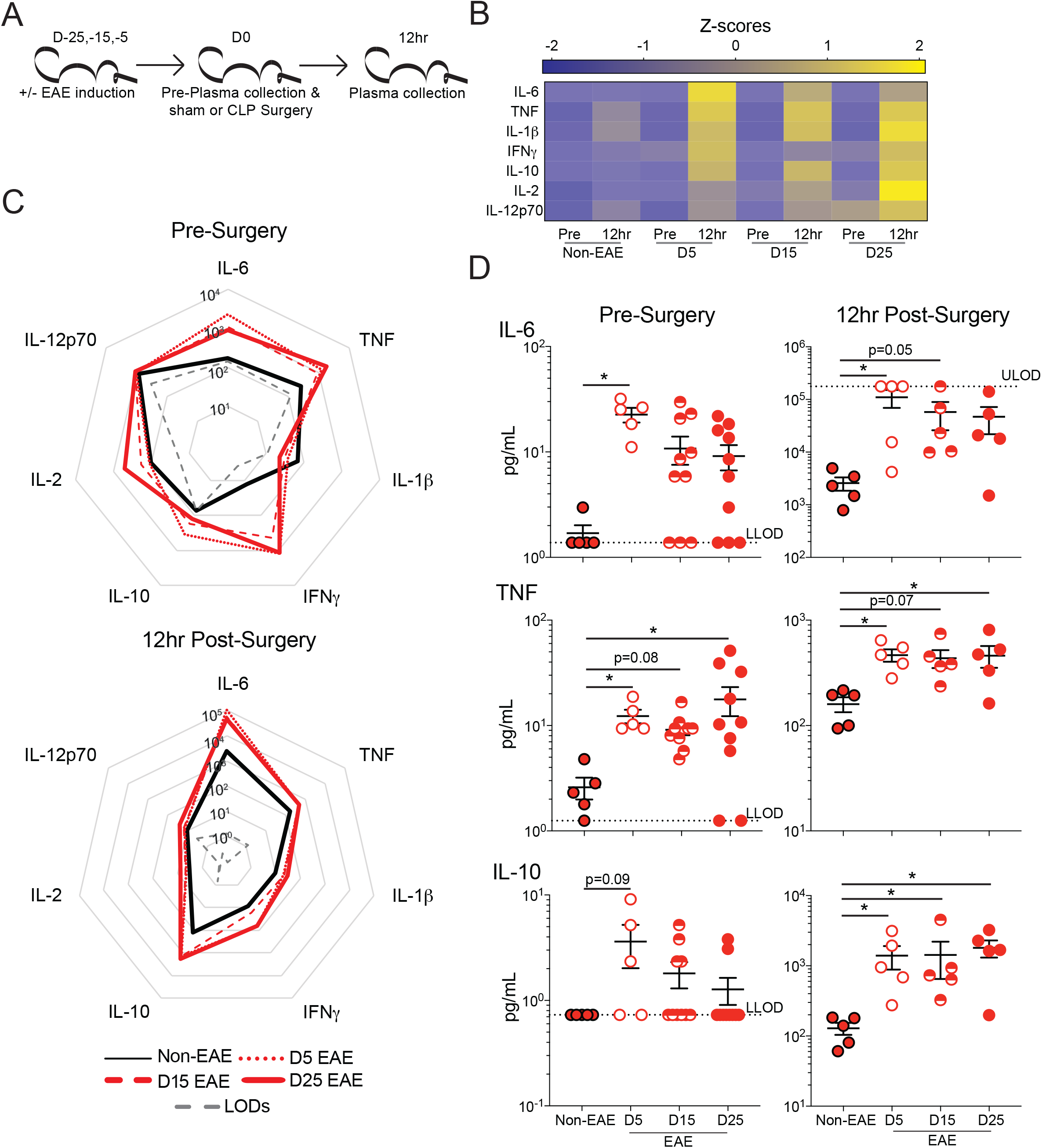
EAE mice have increased inflammation prior to and following sepsis induction. **A)** Experimental Design: C57BL/6 mice were immunized with MOG_35-55_ to induce EAE at day −25, −15, or −5 prior to CLP surgery, age matched non-immunized (non-EAE) underwent CLP surgery at the same time. Plasma was collected prior to surgery and 12 hrs post-surgery. **(B)** Heatmap of normalized plasma IL-6, TNF, IL-1β, IFNγ, IL-10, IL-2, and IL-12p70 concentrations in non-EAE, D5 EAE, D15 EAE, and D25 EAE mice prior to and 12 hrs post-CLP surgery. **(C)** Radar plots of plasma IL-6, TNF, IL-1β, IFNγ, IL-10, IL-2, and IL-12p70 in non-EAE mice (black line), D5 (dotted red line), D15 (dashed red line), and D25 EAE mice (solid red line) prior to (top) and 12 hrs post-(bottom) CLP surgery. **(D)** Representative plasma cytokines (top to bottom: IL-6, TNF, IL-10) prior to (left) and 12hrs post-(right) CLP surgery in non-EAE (red circle with black outline), D5 EAE (white circle with red outline), D15 EAE (red semi-circle), and D25 EAE (red circle with red outline) mice. Grey dashed lines indicate the upper (ULOD) and lower (LLOD) limits of detection for the multiplex assay. Samples are combined from 2 independent experiments run on a single multiplex assay with 5-10 mice per group. Error bars represent standard error of the mean. *=p-value<0.05.

These results then led us to question whether there was a quantitative difference in the magnitude of the cytokine storm between survivor and non-survivor mice at D35 post-EAE induction. Thus, plasma IL-6 and IL-10 were interrogated in survivor and non-survivor EAE mice as well as non-EAE mice prior to and 12 hrs after EAE induction (**Figure 3-figure supplement 1**). Indeed, non-survivor mice had an elevated cytokine storm while survivor mice had a similar magnitude of the cytokine storm as non-EAE mice. This finding further illustrates the susceptibility of EAE mice to sepsis-induced mortality is through enhancement of the cytokine storm.

### EAE mice have increased susceptibility to various models of sepsis induction

Given the high susceptibility of EAE mice to fatal CLP-induced sepsis, we sought to extend the applicability of this effect to other models of sepsis induction. Intraperitoneal injection of lipopolysaccharide (LPS) is a well-established model of endotoxemia and sepsis with a highly tunable degree of mortality by modulating the concentration of LPS (Danahy et al., 2016; Dickson and Lehmann, 2019). With this system, a dose of LPS that elicits a robust cytokine storm, but does not elicit mortality in unmanipulated (e.g., non-EAE) mice, was interrogated (Huggins et al., 2019). LPS was injected 15 days post-EAE induction on EAE and non-EAE cohorts, and mortality was monitored throughout with plasma IL-6 evaluated prior to and 12 hrs post-LPS injection (**Figure 4a**). Consistent with prior experiments, EAE mice had a range of disease scores (**Figure 4b**). Importantly, while non-EAE mice exhibited no mortality, as anticipated, EAE mice exhibited rapid and profound mortality recapitulating the observations with CLP (**Figure 4c**). The enhanced mortality of EAE mice was attributable to increased IL-6 following LPS injection (**Figure 4d**), similar to observations with CLP mice. These data demonstrate increased sensitivity to TLR4 stimulation likely contributes to the enhanced mortality among EAE mice.

**Figure 4:**
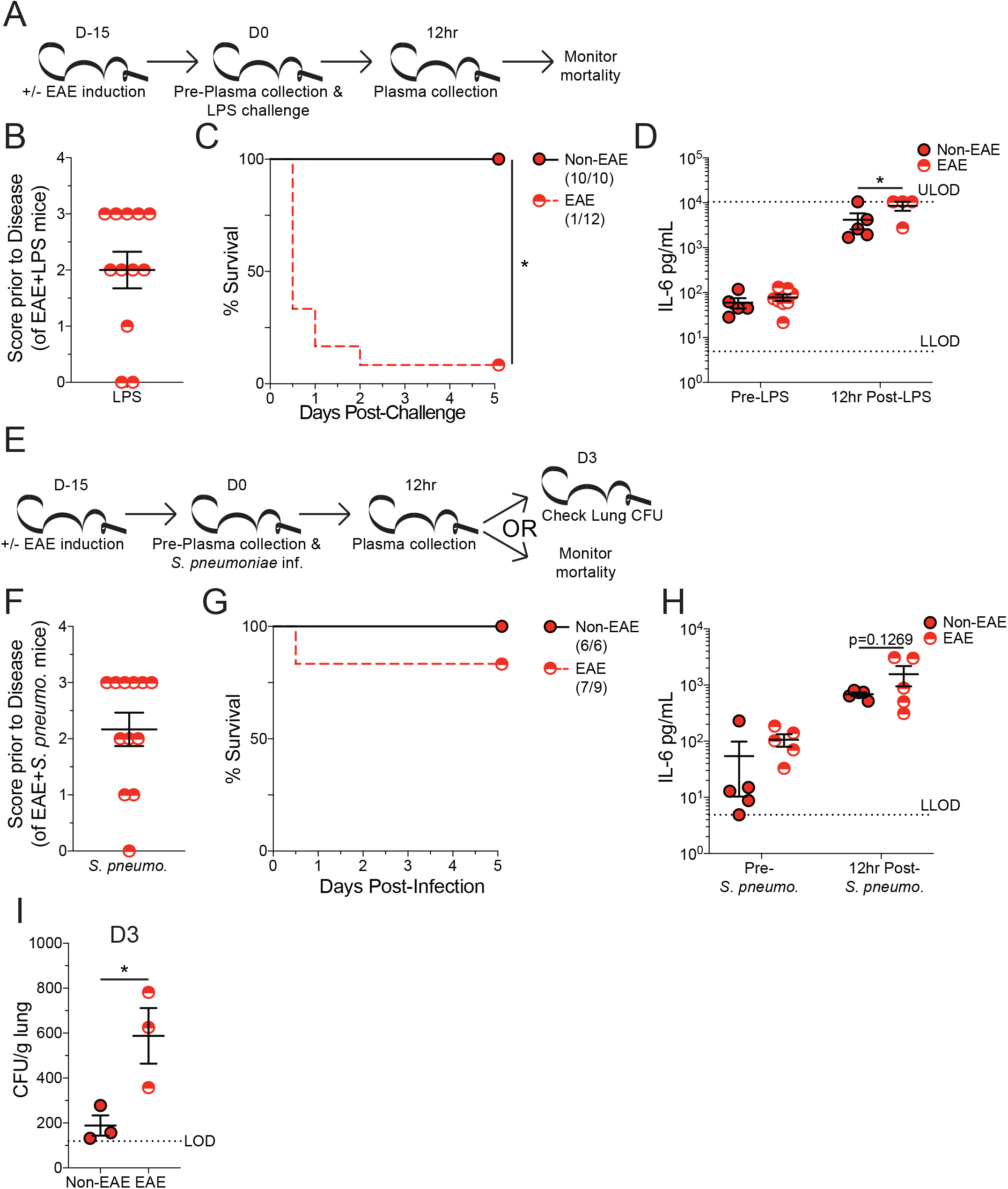
EAE mice have increased susceptibility to multiple sepsis models. **A)** Experimental Design: C57BL/6 mice were immunized with MOG_35-55_ to induce EAE 15 days prior to intraperitoneal LPS injection, age matched non-immunized (non-EAE) received identical injections. Plasma was collected prior to and 12 hrs post-LPS injection. Cohorts were monitored for survival. **(B)** EAE disease scores prior to LPS injection of EAE mice. **(C)** Kaplan-Meier survival curves for non-EAE and D15 EAE mice following LPS injection. **(D)** Plasma IL-6 prior to and 12 hrs post-LPS injection in non-EAE (red circle with black outline) and D15 EAE (red semi-circle). Grey dashed lines indicate the upper (ULOD) and lower (LLOD) limits of detection for IL-6 ELISA. **(E)** Experimental design: C57BL/6 mice were immunized with MOG_35-55_ to induce EAE 15 days prior to intranasal *S. pneumoniae* infection, age matched non-immunized (non-EAE) received identical infections. Plasma was collected prior to and 12 hrs post-LPS injection. 3 mice from each cohort were used for determining lung CFU at 3 days post-infection. Remaining mice in each cohort were monitored forsurvival. **(F)** EAE disease scores prior to *S. pneumoniae* infection of EAE mice. **(G)** Kaplan-Meier survival curves for non-EAE and D15 EAE mice following *S. pneumoniae* infection. **(H)** Plasma IL-6 prior to and 12hrs post-*S. pneumoniae* infection in non-EAE (red circle with black outline) and D15 EAE (red semi-circle). Grey dashed line indicates the lower (LLOD) limits of detection for IL-6 ELISA. **(I)***S. pneumoniae* CFU per gram of lung tissue 3 days after intranasal infection of non-EAE and D15 EAE mice. Dashed line indicates the limit of detection (LOD). Data are from a single experiment with 9-12 mice per group. Error bars represent standard error of the mean. *=p-value<0.05.

Next, we examined the impact of having EAE followed by an intranasal *Streptococcus pneumoniae* (*S. pneumoniae*) infection. *S. pneumoniae* is the most prevalent causative pathogen of community acquired pneumonia, and *S. pneumoniae* models of sepsis have high clinical relevance, as nearly half of all sepsis cases result from this bacterial infection (Brown, 2012). Similar to the LPS endotoxemia model, *S. pneumoniae* infection in this system does not lead to mortality in unmanipulated mice. It does, however, represent a relevant respiratory infection (Bogaert et al., 2004), which are both a common cause of sepsis (Jeganathan et al., 2017) and a frequent complication among MS patients (Harding et al., 2020). Further host ability to control the infection can be assessed by determining the number of colony forming units (CFUs) in the lungs and plasma cytokines to give an indication of the host ability to mount an inflammatory response and clear infection. Utilizing this system, EAE mice and non-EAE controls were intranasally inoculated with *S. pneumoniae* 15 days post-EAE induction. Plasma IL-6 was evaluated prior to and 12 hrs post-*S. pneumoniae* infection. Additionally, lung *S. pneumoniae* CFUs were evaluated in 3 mice from each cohort 3 days after infection while mortality was monitored in the remaining mice (**Figure 4e**). As before, EAE mice exhibited a range of disease severity prior to infection (**Figure 4f**) and some mortality subsequent to the insult (**Figure 4g**), though this mortality was not significantly different from non-EAE control mice. Further, a trending increase in plasma IL-6 was observed from EAE mice 12 hrs post-*S. pneumoniae* infection (**Figure 4h**), in agreement with the prior findings of an elevated inflammatory response in EAE mice challenged with either CLP or LPS. Interestingly, EAE mice also had reduced control of *S. pneumoniae* infection 3 days post-infection, relative to non-EAE mice (**Figure 4i**). These data indicates that despite enhanced inflammation, EAE mice have a dysregulated inflammatory response that has reduced capacity to provide protection to subsequent insult. Thus, the culmination of enhanced inflammatory responses with a reduced capacity to control pathogen insult may set the stage for the enhanced susceptibility of EAE mice, and MS patients, to develop and succumb to septic insults.

Cumulatively, these findings indicate that MS patients are at a higher risk of developing sepsis and ongoing autoimmune reactions lay the groundwork for an exacerbated inflammatory response during septic insult that in turn increases the risk of mortality. This conclusion is relevant to both the identification and management of patient populations that are likely to become septic and at high risk of mortality in the event they become septic. Future work should interrogate the utility of intervention strategies in promoting survival of sepsis and assessments of intervention strategies should account for these highly relevant comorbidities in determining efficacy. Importantly, patients with autoimmunity tend be on immunosuppressive regimens (Tintore et al., 2019; Yong and Kim, 2020), while it is yet unclear what the net result of these interventions are on the development of sepsis, these immunosuppressive regimens will undoubtedly be pertinent to the management of the cytokine storm.

Alternately, it is also relevant to consider the consequences for a patient with autoimmunity who survives a septic insult. This notion is highly related to our previous findings, wherein we observed sepsis-induced immunoparalysis ablated the subsequent development of EAE through the numeric reduction in naïve autoantigen specific CD4 T cells (Jensen et al., 2020). Indeed, sepsis similarly reduces the number and function effector and memory T cells (Cabrera-Perez et al., 2014; Danahy et al., 2017; Duong et al., 2014; Martin et al., 2020; Sjaastad et al., 2020b). Therefore, it is plausible for those individuals that survive to experience a reduction in their autoimmune disease symptoms. Contrastingly, sepsis may also reduce the capacity of suppressor cell populations to mediate their activity and lead to disease exacerbation (Cavassani et al., 2010; Scumpia et al., 2006; Sharma et al., 2015). There are likely multiple complicating factors that dictate whether any such benefit or detriment arises, including the stage of autoimmune disease progression. Such interrogation may lead to enhanced understanding of the sepsis-induced immunoparalysis state or even future therapeutic intervention for MS and autoimmune disease patients.

Finally, it is relevant to consider the observation that clinical disease was not required for the enhancement in mortality among EAE mice. This finding suggests individuals with subclinical or newly developing autoimmunity may be at risk for increased mortality from sepsis. This possibility may be problematic for delineating patient populations with high susceptibility to sepsis-induced mortality as it may not be a recognized complicating factor. Thus, enhanced susceptibility of patient populations to sepsis-induced mortality may be better understood as a result of active inflammatory responses prior to septic insult rather than highly specific comorbidities such as autoimmunity or cancer. These are highly relevant notions when seeking to promote survival and develop future therapeutics.

## Acknowledgements

We thank members of our laboratories and the lab of Dr. Karandikar for technical assistance and helpful discussions.

## Methods

| Key Resources Table |  |  |  |  |
| --- | --- | --- | --- | --- |
| Reagent type (species) or resource | Designation | Source or reference | Identifiers | Additional information |
| strain, strain background<br>(*Mus musculus*) | C57BL6/J | Jackson Laboratory | Stock No: 000664<br>(RRID:IMSR_JAX:000664) |  |
| peptide, recombinant protein | MOG<sub>35-55</sub> | GenScript | SC1208 |  |
| other | CFA containing *M. tuberculosis* H37Ra | Difco | DF3114-33-8 |  |
| peptide, recombinant protein | Pertussis toxin from *Bordetella pertussis* | Sigma-Aldrich | P7208 |  |
| peptide, recombinant protein | LPS (*E. coli* O55:B5) | Sigma | L2880-25MG | 2.5mg/kg of mouse body weight |
| strain, strain background<br>(*Streptococcus pneumoniae*) | *Streptococcus pneumoniae* | Wanke-Jellinek et al. Beneficial Effects of CpG-Oligodeoxynucleotide Treatment on Trauma and Secondary Lung Infection. J Immunol. 196(2):767-77 (2016). |  | Can be acquired through lab contact |
| commercial assay or kit | ProcartaPlex 7-plex | Thermo-Fischer | AB_2575918<br>AB_2575931<br>AB_2575930<br>AB_2575920<br>AB_2575919<br>AB_2575917<br>AB_2575926 | IL-6, TNF, IL-1β, IL-10, IFNγ, IL-2, IL-12p70 |
| antibody | IL-10<br>(JES5-16E3) | Biolegend | AB_315358 | ELISA<br>(1µg/mL) |
| antibody | IL-10<br>(JES5-2A5) | Biolegend | AB_315349 | ELISA<br>(2µg/mL) |
| antibody | IL-6<br>(MP5-32C11) | Biolegend | AB_2233898 | ELISA<br>(1µg/mL) |
| antibody | IL-6<br>(MP5-20F3) | Biolegend | AB_315336 | ELISA<br>(2µg/mL) |
| software,<br>algorithm | GraphPad Prism | GraphPad<br>Prism 8 | Version 8.4.2<br>(464)<br>(RRID:SCR_0<br>02798) |  |

### Retrospective Patient Assessment

TriNetX was utilized to query a limited, deidentified dataset of patients at the University of Iowa admitted between 2008 and 2020. Adult patients (age 18 to 119 years) who had inpatient encounters were queried. Since this period spans the transition from ICD-9 to ICD-10 coding, the TriNetX software uses algorithms to transform ICD-9 codes to ICD-10 codes. Sepsis patients were queried for all ICD-10 codes including “sepsis” in their description utilizing the [or] operator. Multiple sclerosis patients were queried using ICD-10 code group G35 Multiple sclerosis. TriNetX is compliant with the Health Insurance Portability and Accountability Act (HIPAA), the US federal law which protects the privacy and security of healthcare data. TriNetX is certified to the ISO 27001:2013 standard and maintains an Information Security Management System (ISMS) to ensure the protection of the healthcare data it has access to and to meet the requirements of the HIPAA Security Rule. Any data displayed on the TriNetX Platform in aggregate form, or any patient level data provided in a data set generated by the TriNetX Platform, only contains de-identified data as per the de-identification standard defined in Section §164.514(a) of the HIPAA Privacy Rule. The process by which the data is de-identified is attested to through a formal determination by a qualified expert as defined in Section §164.514(b)(1) of the HIPAA Privacy Rule. TriNetX is supported by the Institute for Clinical and Translational Science at the University of Iowa. The Institute for Clinical and Translational Science at the University of Iowa is supported by the National Institutes of Health (NIH) Clinical and Translational Science Award (CTSA) program, grant UL1TR002537. The CTSA program is led by the NIH’s National Center for Advancing Translational Sciences (NCATS). This publication’s contents are solely the responsibility of the authors and do not necessarily represent the official views of the NIH.

### Ethics statement

Experimental procedures using mice were approved by University of Iowa Animal Care and Use Committee under ACURF protocol #6121915 and #9101915. The experiments performed followed Office of Laboratory Animal Welfare guidelines and PHS Policy on Humane Care and Use of Laboratory Animals. Cervical dislocation was used as the euthanasia method of all experimental mice.

### Mice

Inbred C57Bl/6 (B6; Thy1.2/1.2) were purchased from the National Cancer Institute (Frederick, MD) and maintained in the animal facilities at the University of Iowa at the appropriate biosafety level.

### Cecal ligation and puncture (CLP) model of sepsis induction

CLP surgery was performed as previously described (Sjaastad et al., 2020a). Briefly, mice were anesthetized with ketamine/xylazine (University of Iowa, Office of Animal Resources), the abdomen was shaved and disinfected with Betadine (Purdue Products), and a midline incision was made. The distal third of the cecum was ligated with Perma-Hand Silk (Ethicon), punctured once using a 25-gauge needle, and a small amount of fecal matter extruded. The cecum was returned to abdomen, the peritoneum was closed with 641G Perma-Hand Silk (Ethicon), and skin sealed using surgical Vetbond (3M). Following surgery, 1 mL PBS was administered s.c. to provide post-surgery fluid resuscitation. Lidocaine was administered at the incision site, and flunixin meglumine (Phoenix) was administered for postoperative analgesia. This procedure created a septic state characterized by loss of appetite and body weight, ruffled hair, shivering, diarrhea, and/or periorbital exudates with 0–10% mortality rate. Sham mice underwent identical surgery excluding cecal ligation and puncture.

### LPS Endotoxemia induction

Mice received a single intraperitoneal injection of LPS-EB from *E. coli* O55:B5 (2.5 mg/kg body weight; Sigma), as previously described (Huggins et al., 2019).

### *Streptococcus pneumoniae* infection

Streptococcus was grown in brain heart infusion (BHI) broth then pelleted by centrifugation. Pellet was washed three times and diluted to a target absorbance of 0.1 using PBS, as measured by ABS_600_. Mice were anesthetized with ketamine/xylazine and received 40μL of *Streptococcus pneumoniae* by intranasal inoculation. Infectious dose was confirmed by plating inoculum (1.5×10^6^ CFU/ mouse) on BHI plates.

CFU per gram of lung was determined by sacrificing mice and weighing the lungs. Lungs were mechanically homogenized in 1mL of PBS. 20μL of homogenate on BHI plates in duplicate.

### EAE Disease Induction and Evaluation

EAE was induced and evaluated as shown previously (Mangalam et al., 2009). Briefly, mice were immunized s.c. on day 0 on the left and right flank with 100 μg of MOG_35-55_ emulsified in Complete Freund’s Adjuvant followed by 80 ng of pertussis toxin (PTX) i.p. on days 0 and 2. Disease severity was scored as follows: 0, no clinical symptoms; 1, loss of tail tonicity; 2, hind limb weakness; 3, hind limb paralysis; 4, fore limb weakness; 5, moribund or death.

### Cytokine Analysis

Multiplex cytokine analysis was performed via Thermo-Fischer ProcartaPlex 7-plex according to the manufacturer’s instructions for plasma cytokine analysis. Multiplex was analyzed on BioRad Bio-Plex (Luminex 200) analyzer in the University of Iowa Flow Cytometry core facility.

IL-6 and IL-10 ELISAs (ELISA MAX Deluxe Set, Biolegend) were performed according to the manufacturer’s instructions.

### Statistical Analysis

Unless stated otherwise data were analyzed using Prism 8 software (GraphPad) using two-tailed Student t-test (for 2 individual groups, if variance was unequal variance then Mann-Whitney U test), one-way ANOVA with Bonferroni post-hoc test (for >2 individual groups, if variance was unequal variance then Kruskal-Wallis with Dunn’s post-hoc test was used), two-way ANOVA (for multiparametric analysis of 2 or more individual groups, pairing was used for samples that came from the same animal) with a confidence interval of >95% to determine significance (*p < 0.05). Log-rank (Mantel-Cox) curve comparisons was used to determine significant difference in time to disease EAE disease onset (*p < 0.05). Data are presented as standard error of the mean.

## Source data

Figure 1-source data 1

Source data for Figure 1.

Figure 2-source data

Source data for Figure 2.

Figure 3-source data

Source data for Figure 3.

Figure 4-source data 1

Source data for Figure 4B, C.

Figure 4-source data 2

Source data for Figure 4D.

Figure 4-source data 3

Source data for Figure 4F, G.

Figure 4-source data 4

Source data for Figure 4H.

Figure 4-source data 5

Source data for Figure 4I.

Figure 3-figure supplement 1-source data 1

Source data for Figure 3-figure supplement 1 A, B.

Figure 3-figure supplement 1-source data 2

Source data for Figure 3-figure supplement 1 C, D.

Table 1-source data

Source data for Table 1.

---

**Figure 3-figure supplement 1:**
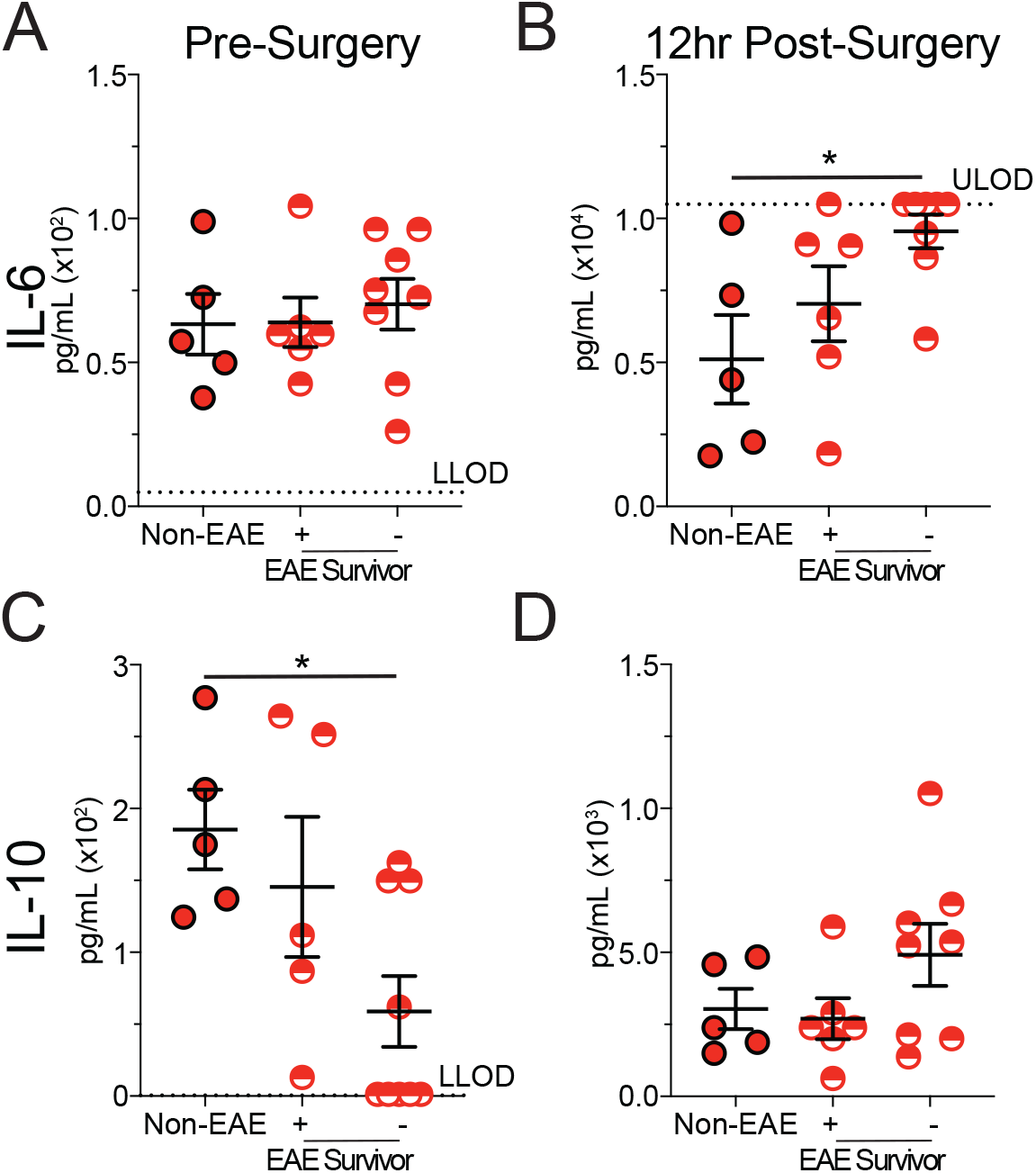
Mortality in EAE mice is associated with elevated inflammation. C57BL/6 mice were immunized with MOG_35-55_ to induce EAE. EAE and age matched non-immunized (non-EAE) mice underwent CLP 35 days after EAE induction. Plasma cytokines were assessed prior to and 12 hrs post-CLP surgery in non-EAE, EAE mice that survived CLP-induced sepsis, and EAE mice that succumbed to CLP-induced sepsis. Plasma IL-6 **(A, B)** and IL-10 (**C, D)** prior to **(A, C)** and 12 hrs post-**(B, D)** CLP surgery in non-EAE, EAE mice that survived CLP-induced sepsis, and EAE mice that succumbed to CLP-induced sepsis. Grey dashed lines indicate the upper (ULOD) and lower (LLOD) limits of detection for the respective ELISA plate. Samples are combined from 2 independent experiments run on single ELISA plates with 5-8 mice per group. Error bars represent standard error of the mean. *=p-value<0.05.

